# Intrinsically disordered domain of kinesin-3 Kif14 enables unique functional diversity

**DOI:** 10.1101/2020.01.30.926501

**Authors:** Ilia Zhernov, Stefan Diez, Marcus Braun, Zdenek Lansky

## Abstract

In addition to their force-generating motor domains, kinesin motor proteins feature various accessory domains enabling them to fulfil a variety of functions in the cell. Human kinesin-3, Kif14, localizes to the midbody of the mitotic spindle and is involved in the progression of cytokinesis. The specific motor properties enabling Kif14’s cellular functions, however, remain unknown. Here, we show in vitro that it is the intrinsically disordered N-terminal domain of Kif14 that enables unique functional diversity of the motor. Using single molecule TIRF microscopy we observed that the presence of the disordered domain i) increased the Kif14 run-length by an order of magnitude, rendering the motor super-processive and enabling the motor to pass through highly crowded microtubule areas shielded by cohesive layers of microtubule-associated protein tau, which blocks less processive motors ii) enabled robust, autonomous Kif14 tracking of growing microtubule tips, independent of microtubule end-binding (EB) proteins and iii) enabled Kif14 to crosslink parallel microtubules and to drive the relative sliding of antiparallel ones. We explain these features of Kif14 by the observed increased affinity of the disordered domain for GTP-like tubulin and the observed diffusible interaction of the disordered domain with the microtubule lattice. We hypothesize that the disordered domain tethers the motor domain to the microtubule forming a diffusible foothold. We suggest that the intrinsically disordered N-terminal anchoring domain of Kif14 is a regulatory hub supporting the various cellular functions of Kif14 by tuning the motor’s interaction with microtubules.

## INTRODUCTION

Molecular motors associating with cytoskeletal filaments typically consist of mechanochemically functional enzymatic motor-domains propelling directed translocation, accompanied by a modular set of task-specific domains or accessory proteins. Of the three classes of mechanochemically active motors, kinesins, myosins and dyneins, the cytoplasmic dynein complex, while itself remaining unaltered, is regulated by associated proteins and complexes, for example, the dynactin complex and BicD [1, 2]. Following a different evolutionary strategy, both the myosins and kinesins differentiated into large families of distinct and diverse proteins, which, alongside the conserved motor domains, each comprise different functional domains [3, 4]. These accessory functional domains enable the motors to fulfill various roles in the cell, as for example sliding microtubules relative to each other [5–7] or autonomously crosslinking microtubules and actin filaments [8, 9], enabling EB1-dependent tip-tracking of growing microtubule ends [10] or assembly [11], and providing interaction sites for binding to other microtubule-associated proteins [12–14].

Motors of the kinesin-3 are involved in a large variety of cellular processes, such as intracellular cargo transport or mitosis [15]. Human kinesin-3, Kif14 enriches in the spindle midbody and the central spindle during cytokinesis [16, 17], where it interacts with protein regulator of cytokinesis 1 (PRC1) and is essential for localizing citron kinase to the mitotic spindle [16, 17]. Depletion of Kif14 disrupts cell cycle progression and induces cytokinesis failure, resulting in multinucleation and apoptosis [16, 17]. Kif14 contributes to the establishment and maintenance of the cytokinetic furrow through interaction with supervillin [18] and Kif14 overexpression is associated with various cancers [19]. Kif14 has been also implicated in brain development, as mutations of Kif14 have been described in patients with primary microcephaly [20–22], brain malformation [23], and Meckel syndrome 12 [24]. Molecular mechanisms enabling Kif14 to fulfill its multiple roles are currently unknown. However, Kif14, in addition to the structurally and biochemically well described motor domain [25], uniquely features at its N-terminus an extended intrinsically disordered protein domain. Apart from being involved in the Kif14 interaction with PRC1 and F-actin [17, 26], the functionality of this domain is largely unknown.

We here demonstrate in vitro that this intrinsically disordered protein domain of the kinesin-3 Kif14 serves as a diffusible anchor, tethering the protein to the microtubule lattice, promoting Kif14 super-processivity, autonomous tip tracking, stable crosslinking of parallel microtubules and relative sliding of antiparallel ones. This suggests diffusible tethering of the Kif14 motor domains as means to promote long-range motility, and, in dividing cells, after ensuring the localization of Kif14 to the spindle midbody, the stabilization of the midbody, increasing the robustness of the central spindle during cytokinesis.

## RESULTS

### 1. Intrinsically disordered domain promotes Kif14 processivity

Apart from the motor domain (aa 355-709), a coiled-coil (aa 709-800), a forkhead-associated domain (aa 800-900), and a stalk with a tail (aa 900-1648), Kif14 possesses an unusually long stretch of amino acids located N-terminally (aa 1-355) of the motor domain. Bioinformatic analysis revealed this domain as intrinsically disordered and positively charged with an isoelectric point of 9.63 (Fig. S1,S2). The positive charge of the disordered domain suggests a possible direct interaction with microtubules. To determine the influence of the disordered domain on the Kif14 motor activity we first purified a recombinant GFP-tagged minimal dimerizable construct, Kif14(768)-eGFP (aa 1-768) (Fig. S3), comprising the disordered domain, the motor domain and the coiled coil (Fig. S1b). We then immobilized GMPCPP-stabilized microtubules on the surface of the flow chamber by anti-β-tubulin antibodies and visualized the Kif14(768)-eGFP interaction with microtubules using TIRF microscopy (Fig. 1a). Imaging the positions of single-molecules we observed processive and diffusive Kif14(768)-eGFP movement, with diffusive molecules comprising the vast majority (89 %) (Fig. 1b). Each individual encounter of Kif14(768)-eGFP with a microtubule was either diffusive or processive, with the molecules never switching between the two modes of motion (Fig. 1b). Processive Kif14(768)-eGFP moved with velocity of 153 ± 51 nm/s (mean ± SD, n = 247 molecules) (Fig. 1e) and exhibited median run-length of 7.32 µm (95% confidence interval, CI_95_ (5.5, 8.75) μm, n = 247 molecules) (Fig. 1d), similar to other superprocessive motors, as e.g. Kip3 [27, 28]. Like Kip3 [27, 28], Kif14(768)-eGFP also accumulated at the ends of the microtubules (Fig. S4a,c; movie 1).

**Fig. 1.**
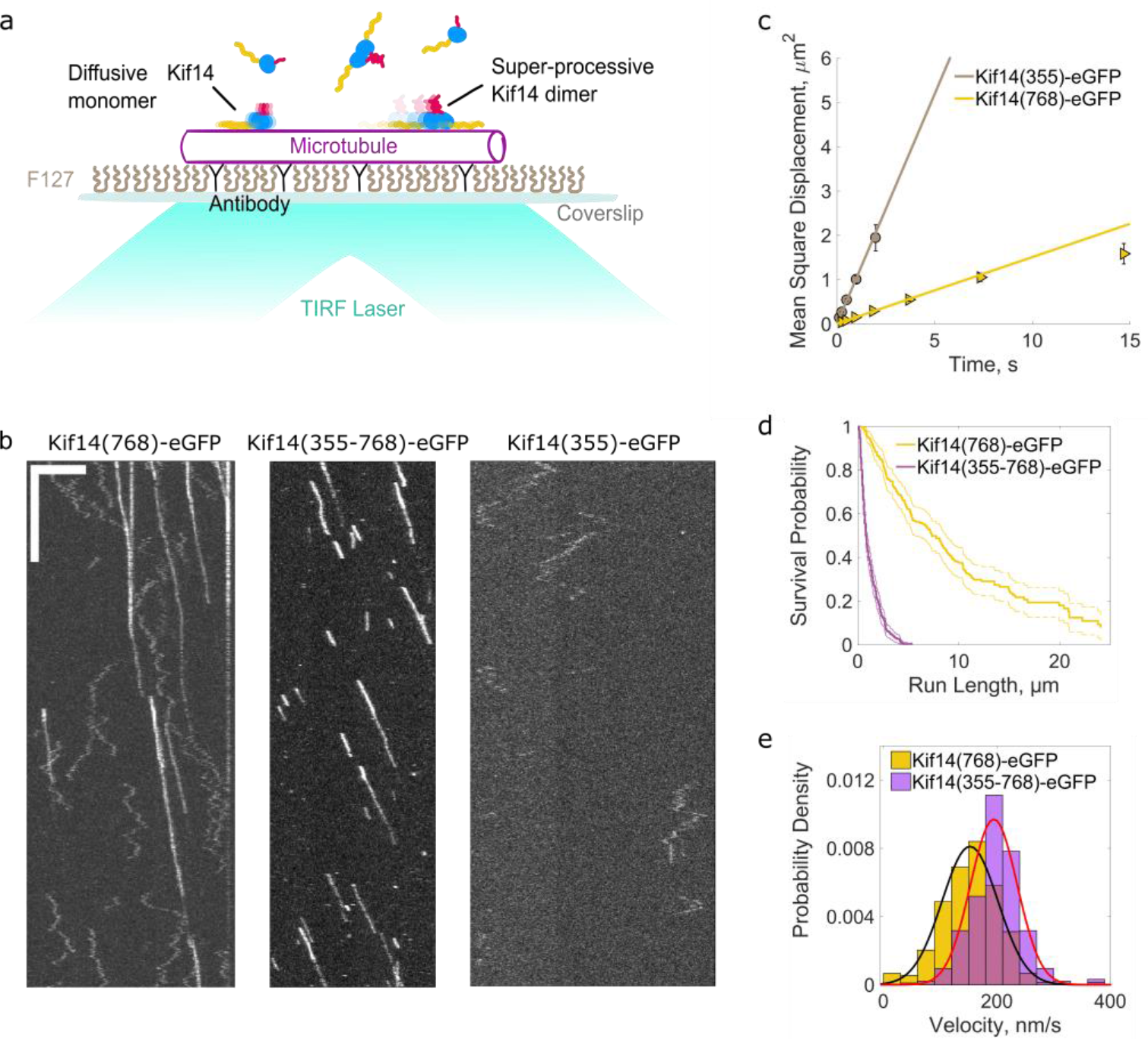
Kif14 intrinsically disordered domain promotes processivity and enables diffusion along microtubules. **(a)** Schematics of a Kif14 stepping assay. **(b)** Kymographs representing motion of Kif14(768)-eGFP, Kif14(355-768)-eGFP and Kif14(355)-eGFP on the microtubule (unlabeled). Horizontal scale bar 5 μm; vertical bar 20 s. **(c)** Mean square displacement of Kif14 constructs: Kif14(355)-eGFP (diffusion constant, D = 0.5180 ± 0.0415 μm^2^/s, n = 142) and Kif14(768)-eGFP (diffusion constant, D = 0.0751 ± 0.0066 μm^2^/s, mean ± 95 % confidence bounds, n = 234). Lines indicate linear fits to the MSD data points, error bars indicate standard deviation. **(d)** Survival probability of run lengths of Kif14(768)-eGFP (yellow) and Kif14(355-768)-eGFP (magenta). Dotted lines indicate 95 % confidence bounds. **(e)** Distribution of velocities of Kif14(768)-eGFP (yellow, 153 ± 51 nm/s, n = 247 molecules) and Kif14(355-768)-eGFP (magenta, 198 ± 49 nm/s, mean ± SD, n = 315 molecules). Black and red line indicate Gaussian fits to the histograms.

We next generated dimerizable motor domain construct lacking the disordered domain, Kif14(355-768)-eGFP and tested its motility. All molecules of this construct moved processively on microtubules (none of them moved diffusively), with decreased median run-length of 0.77 µm (95% confidence interval, CI_95_ (0.69, 0.96) μm, n = 315 molecules) (Fig. 1d). This construct moved significantly faster than Kif14(768)-eGFP (Fig. 1e) with velocity of 198 ± 49 nm/s (mean ± SD, n = 315 molecules), suggesting that the disordered domains’ interaction with the microtubule surface induced drag on the movement of the molecular motor. In accordance with its lower processivity, we observed no end-accumulation of Kif14(355-768)-eGFP (Fig. S4b,d, movie 2). These observations indicate that the Kif14 disordered domain supports the diffusible motion of Kif14 along microtubules and enhances its processivity.

We next tested if the disordered domain was sufficient to support diffusion on microtubules. To this aim we generated a minimal Kif14(355)-eGFP construct, which comprises only the disordered domain. We found that Kif14(355)-eGFP binds-to and diffuses-along microtubules (Fig. 1b) with about 7-fold increase in diffusion constant as compared to Kif14(768)-eGFP (Fig. 1c), indicating that the motor domain influences the diffusive behavior.

Taken together, these experiments show that Kif14(768)-eGFP exhibits two modes of motion on the microtubules, either diffusive or super-processive. We propose, the intrinsically disordered domain anchors the motor domain to the microtubule, and either enables its diffusion, or, reducing the kinesins unbinding rate and increasing its run length, renders the motor super-processive.

### 2. Diffusion and stepping require Kif14 to be monomeric or dimeric, respectively

To disentangle the relation between the Kif14 diffusive and super-processive populations, we first generated an intrinsically monomeric Kif14 construct – Kif14(709)-eGFP, comprising the Kif14 diffusible anchoring domain and the motor domain, but lacking the coiled-coil dimerization region (Fig. S3). The constitutively monomeric Kif14(709)-eGFP exhibited purely diffusive motion, with a diffusion constant very similar to Kif14(768)-eGFP (Fig. 2a; Fig. S5a,b). To determine the oligomerization state of the diffusible and processive populations of the dimerizable Kif14(768)-eGFP, we compared their fluorescent intensities (Fig. 2b) with the fluorescent intensities of the constitutively dimeric kinesin-1 Kif5b-GFP (Methods). Fluorescence intensities of the dimeric Kif5b-GFP molecules formed a bimodal gaussian distribution with a main peak comprising 60 % of the imaged molecules, which we attribute to the signal of two GFPs, and a lesser peak, comprising 40 % of the molecules at about half the intensity of the main population (Fig. 2c). The dimmer population, attributed to single GFP signal, is likely comprised of Kif5b molecules harboring a non-fluorescent eGFP due to misfolding, incomplete maturation of GFP or bleaching events occurring before data acquisition. The intensities of diffusive Kif14(768)-eGFP molecules formed a monomodal distribution with a mean value around the determined intensity of one GFP, indicating that diffusive Kif14(768)-eGFP molecules were monomers (Fig. 2d). The super-processive molecules observed in the same experiment, showed a bimodal distribution as the dimeric Kif5b-GFP (Fig. 2e), indicating that the super-processive Kif14(768)-eGFP molecules are dimers. We thus conclude that Kif14(768)-eGFP molecules exist in two forms, either as diffusible monomers or super-processive dimers. The absence of diffusible dimers suggests that the mode of Kif14(768)-eGFP interaction with microtubules is determined, via coiled coil dimerization.

**Fig. 2.**
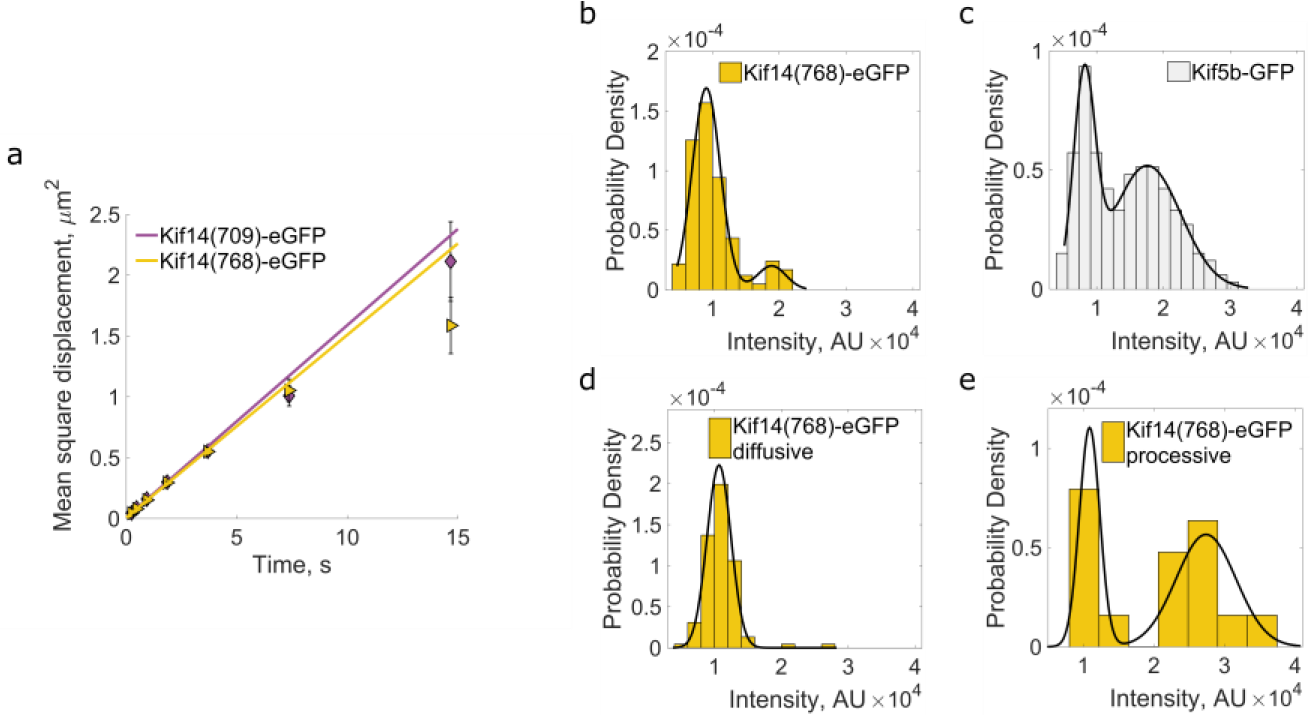
Diffusion and stepping require Kif14 to be monomeric or dimeric, respectively. **(a)** Mean square displacement of Kif14 constructs: Kif14(709)-eGFP (D = 0.0790 ± 0.0069 μm^2^/s, n = 434 molecules) and Kif14(768)-eGFP (D = 0.0751 ± 0.0066 μm^2^/s, mean ± 95 % confidence bounds, n = 234 molecules). Lines indicate linear fits to the MSD data, error bars indicate standard deviation. **(b-e)** GFP signal intensity distribution of: (b) Kif14(768)-eGFP molecules in nucleotide-free state, (c) Kif5b-GFP molecules in nucleotide-free state, (d) Kif14(768)-eGFP diffusive molecules in ATP-containing buffer, (e) Kif14(768)-eGFP processive molecules in ATP-containing buffer. Black lines indicate Gaussian fits to the histograms.

### 3. Enhanced Kif14 processivity enables stepping in crowded environments

Enhanced processivity enhances kinesin motility in crowded environments [29], for example the super processive kinesin-8 Kip3 molecules are able to move through islands of densely-packed tau molecules enveloping the microtubule surface, which hinder the motility of the less processive kinesin-1 [30]. To test if anchoring via the disordered domain might enhance the processivity of Kif14 stepping under such extreme crowding conditions, we formed tau cohesive islands and observed movement of Kif14 molecules within and outside of the islands. We first incubated microtubules immobilized on the surface of the coverslips in the presence of tau-mCherry until tau islands of steady sizes assembled. Next, we infused a mixture of tau-mCherry with Kif14(768)-eGFP or Kif14(355-768)-eGFP in the channel. We observed both Kif14(768)-eGFP and Kif14(355-768)-eGFP binding preferentially to the non-island regions on the microtubules (Fig. 3a,b). Within tau islands Kif14(768)-eGFP, including the anchoring domain, showed significantly higher run lengths than the dimerizable motor-only Kif14(355-768)-eGFP, which does not feature the anchoring domain (Fig. 3c; movie 3, movie 4). Kif14(768)-eGFP often successfully traversed the islands, while Kif14(355-768)-eGFP did not. Thus, we argue that the intrinsically disordered domain, by serving as diffusible anchor that increases the Kif14 processivity, allows Kif14 to move efficiently in crowded environments – as exemplified by its ability to traverse microtubules regions enveloped in cohesive tau islands.

**Fig. 3.**
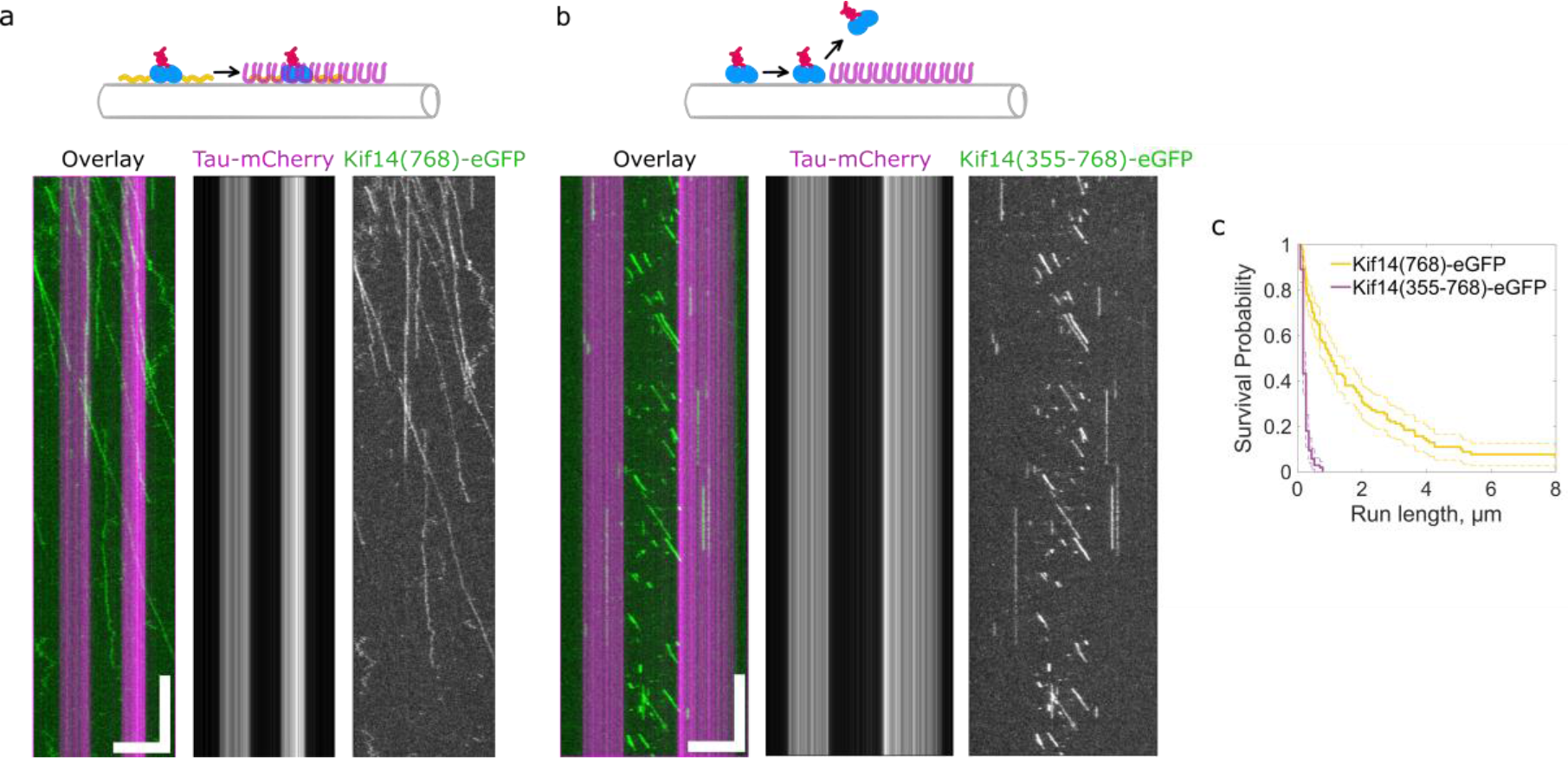
Enhanced Kif14 processivity enables stepping in cohesive tau islands. **(a)** Kymographs representing motion of Kif14(768)-eGFP (green) on the microtubule (unlabeled) covered with tau islands (magenta). Horizontal scale bar 5 μm; vertical bar 2 min. **(b)** Kymographs representing motion of Kif14(355-768)-eGFP (green) on the microtubule (unlabeled) covered with tau islands (magenta). Horizontal scale bar 5 μm; vertical bar 2 min. **(c)** Survival probability plot of run lengths of Kif14(768)-eGFP (yellow) and Kif14(355-768)-eGFP (magenta) in tau islands. Dotted lines indicate 99 % confidence bounds. Median run-length is 1.01 µm, 95% confidence interval (0.69, 1.41) µm, (n = 156 molecules) for Kif14(768)-eGFP and 0.16 µm, 95% confidence interval (0.13, 0.17) µm, (n = 111 molecules) for Kif14(355-768)-eGFP.

### 4. Kif14 autonomously tracks dynamically growing microtubule tips

Super-processive motors of the kinesin-8 family, when forming traffic jams at microtubule plus-ends, are known to induce microtubule depolymerization [27, 28]. We thus wondered if super-processive Kif14 traffic jams accumulation at microtubule plus-ends, analogously, might have a similar effect. To explore this possibility, we employed microtubule dynamics assay. In the presence or absence of different Kif14 constructs, we introduced 20 μM unlabeled porcine tubulin in dynamic buffer to GMPCPP-stabilized microtubule seeds immobilized on the coverslip and visualized the process by TIRF and interference reflection microscopy [31]. Unexpectedly, we found that Kif14(768)-eGFP strongly associated with GMPCPP-stabilized microtubule seeds and growing microtubule tips, while it had a 3-7 fold lower affinity for the GDP-tubulin lattice of dynamically formed microtubules (Fig. 4a-d, movie 5). The indiscriminate association of Kif14(768)-eGFP with growing plus and minus tips indicates that Kif14 might not require super-processivity to localize to growing microtubule tips. To test this, we employed in the dynamics assay Kif14(709)-eGFP comprising the anchoring and the motor domains, but not the coiled coil region and thus being incapable of any processive movement. Similarly to Kif14(768)-eGFP, Kif14(709)-eGFP localized to the seed and the growing tips, indicating that Kif14 tip tracking is mediated by direct preferential binding to the GTP-like tubulin lattice rather than by successive accumulation after super-processive movement along the microtubules (Fig. S6a-c; movie 6). To determine if the anchoring domain, is involved in Kif14 tip tracking and recognition of the GTP-tubulin lattice, we recapitulated dynamic assay using Kif14(355)-eGFP, comprising only the anchoring domain and Kif14(355-768)-eGFP the dimeric motor domains lacking the anchoring domains. Intriguingly, both constructs associate strongly with the microtubule seed, yet less efficiently than Kif14(768)-eGFP or Kif14(709)-eGFP (Fig. S6d-i; movie 7, movie 8), indicating synergy between two autonomously tip-tracking domains, the Kif14 anchoring domain and the motor domain. Surprisingly, Kif14(768)-eGFP tip-tracking did not alter any of the dynamic instability parameters (Fig. 4e-h). Collectively, these observations demonstrate that the anchoring domain and the motor domain of Kif14 both preferentially bind GTP tubulin lattice over GDP-tubulin lattice and thus synergistically enable Kif14 to autonomously track the tips of polymerizing microtubules.

**Fig. 4.**
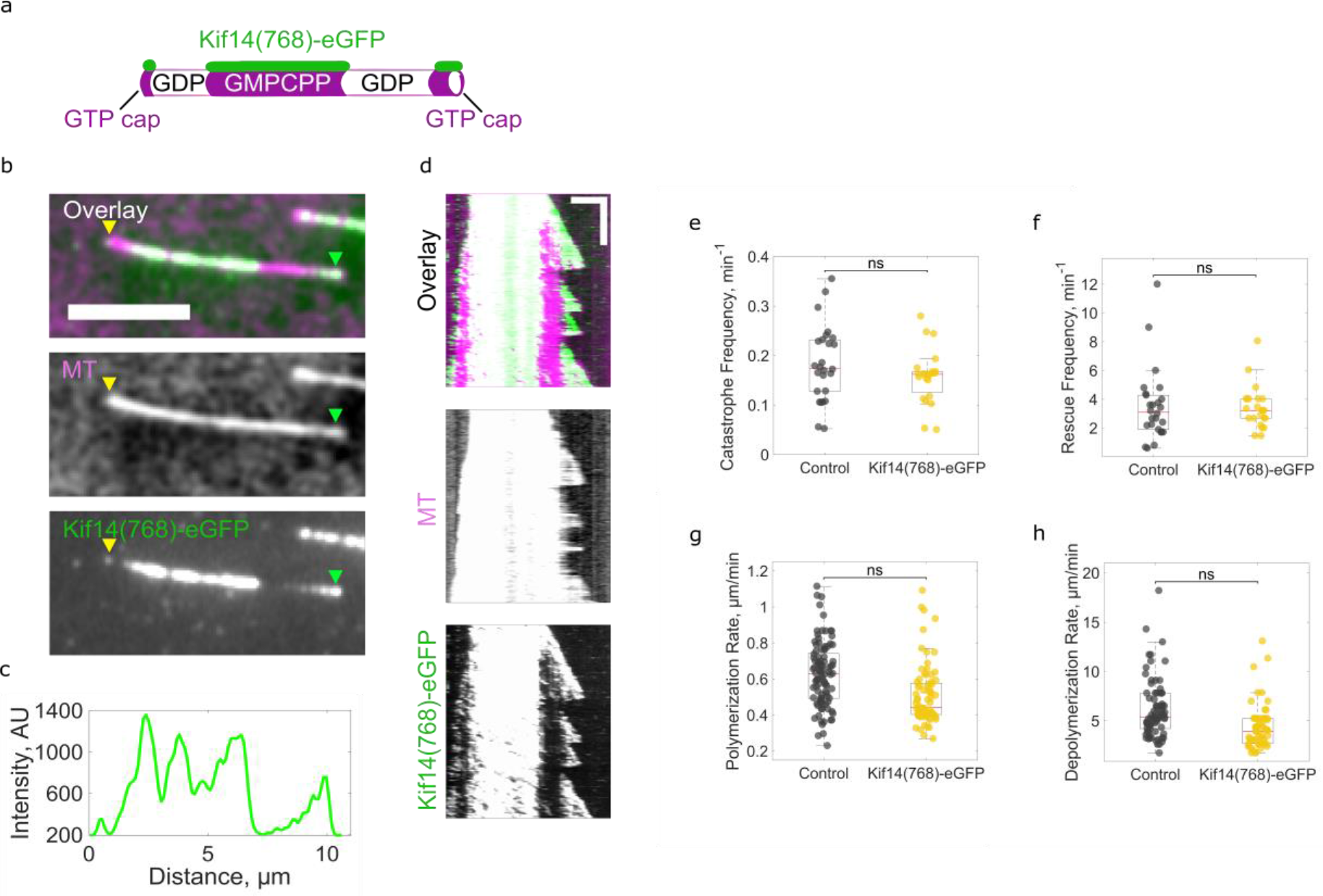
Kif14 preferentially binds GTP-bound microtubule lattice and autonomously tracks dynamically growing microtubule tips. **(a)** Schematics of dynamic microtubule assay. **(b)** Snapshots of a dynamic microtubule (magenta) in the presence of 10 nM Kif14(768)-eGFP (green); green and yellow arrowheads indicate plus and minus microtubule tips respectively. **(c)** Intensity distribution of the GFP signal along the microtubules in (b). **(d)** Kymographs representing 10 nM Kif14(768)-eGFP (green) interaction with the polymerizing microtubule (magenta). Horizontal scale bar 3 μm; vertical bar 5 min. **(e-h)** Dynamic instability parameters, namely (e) catastrophe frequency, (f) rescue frequency, (g) polymerization rate, (h) depolymerization rate in the presence or absence of 10 nM Kif14(768)-eGFP (ns – not significant, determined by Student t test).

### 5. Kif14 crosslinks and slides microtubules

Kif14 is a crucial component of the midbody, where it orchestrates cytokinesis together with other microtubule-binding proteins. A number of mitotic kinesins crosslink and slide microtubules along each other via their multiple binding domains. Given the two microtubule-binding domains, the anchoring domain and the motor domain, Kif14 might be involved in microtubule-microtubule crosslinking and sliding. We thus tested if Kif14 is capable of microtubule sliding assay by immobilizing biotinylated template microtubules on the coverslip surface via anti-biotin antibodies and subsequently flushing-in a mixture of rhodamine labeled transport microtubules and 20 nM Kif14(768)-eGFP or 300 nM Kif14(355-768)-eGFP in motility buffer (Fig. 5a). In the presence of Kif14(768)-eGFP we observed robust landing of transport microtubules on the template microtubules. 40% of transport microtubules that landed on template microtubules showed directional sliding, while 60% were statically crosslinked (Fig. 5b-c; movie 9). Sliding transport microtubules moved towards the plus tips of template microtubules with their minus end leading, as assessed by the Kif14(768)-eGFP end accumulation, indicating anti-parallel arrangement of the template and transport microtubules (Fig. 5b). Conversely, parallel arrangement leaded to static crosslinking. These results suggest a directional sliding mechanism, similar to other mitotic kinesins [5–7, 32]. In the presence of Kif14(355-768)-eGFP we observed transport microtubules landing on template microtubules only very sporadically, indicating the importance of the diffusible anchor domain for the process (Fig. 5a, Fig. S7). By contrast, Kif14(355)-eGFP, comprising the diffusible anchor only, readily crosslinked microtubules, which then diffused relative to each other, but did not slide directionally (Fig. S8), demonstrating that the Kif14 anchoring domain is sufficient and the motor domain is dispensable for microtubule crosslinking. Furthermore, as Kif14(355)-eGFP lacks the coiled coil domain, this data also demonstrates that individual anchoring domains are capable of crosslinking microtubules, and that dimerization is not required. Taken together, these data show that the Kif14 anchoring domain efficiently crosslinks microtubules and, cooperating with the motor domain, allows Kif14 to propel relative microtubule sliding, in an orientation dependent manner.

**Fig. 5.**
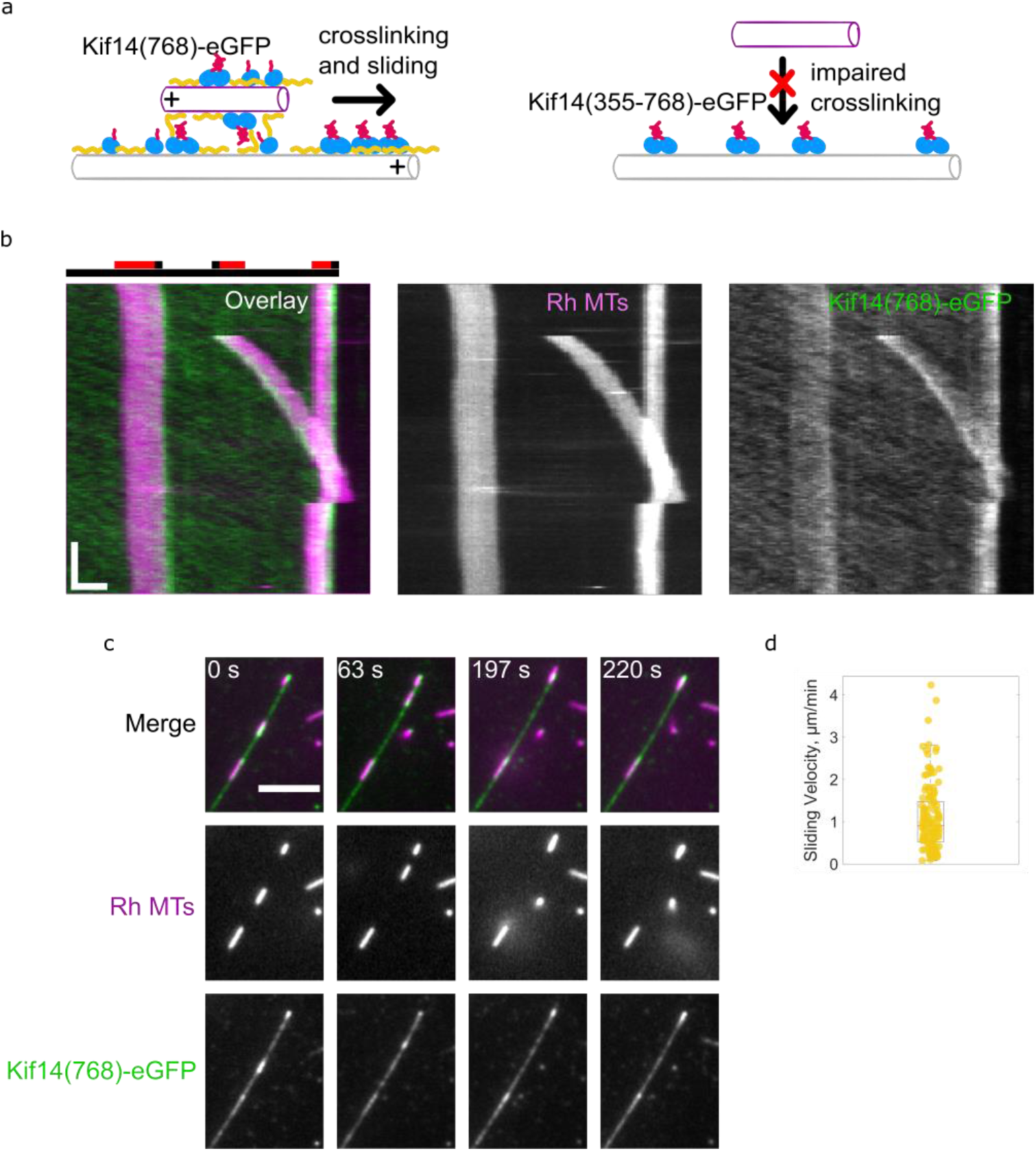
Kif14 crosslinks and slides microtubules. **(a)** Schematics of microtubule sliding by Kif14(768)-eGFP. **(b)** Kymographs representing a rhodamine-labeled transport microtubule (magenta) moved by Kif14(768)-eGFP (green) along a template microtubule (unlabeled). The schematic diagram illustrates microtubule orientations and positions at the start of the experiment (template microtubule is gray and transport microtubules are red; black marks indicate plus-ends). The short transport microtubule on the right presumably has a parallel orientation and therefore is not moved by Kif14(768)-eGFP. **(c)** Time-lapse fluorescence micrographs of a rhodamine-labeled transport microtubule (magenta) sliding along a template microtubule (unlabeled) in the presence of 20 nM Kif14(768)-eGFP. **(d)** Velocity of microtubule-microtubule sliding by Kif14(768)-eGFP: 1.1 ± 0.81 µm/min (mean ± SD, n = 122 molecules).

## DISCUSSION

Similarly to crosslinkers of the kinesin-5 and −14 family members [6–9, 33–37], kinesin-3, Kif14, can slide antiparallel microtubules relative to each other, while statically crosslinking parallel ones, consistent with the tug-of-war mechanism proposed earlier [5, 7]. Intriguingly, unlike kinesin-14, crosslinking by Kif14 does not require the motor domain, showing that the Kif14 disordered anchoring domain can interact with two adjacent microtubules simultaneously and thereby establish diffusible crosslinks. Spindle midzone microtubules are crosslinked mainly by PRC1 at an average distance ranging from 19 to 35 nm [38, 39]. The anchoring domain stretches over 350 amino acids, is intrinsically disordered, and may thus, with a contour length of 122 nm, comfortably span between adjacent microtubules in a bundle. Taken together, these results show that Kif14 may be added to the distinct set of mitotic kinesins actively involved in crosslinking and sliding microtubules and might be thus involved in establishing and maintaining the three-dimensional shape of the microtubule spindle during cell division.

Our data reveal preferred binding of the Kif14 anchoring domain to microtubule domains consisting of tubulin in the GTP-like state. Despite being intrinsically disordered, the Kif14 diffusible anchoring domain thus seems to bind in a distinct conformation to the microtubule surface, possibly analogously to the R4 microtubule binding repeats of tau [40]. Similar to the anchoring domain also Kif14 motor domains preferentially associates with GTP-like tubulin lattice. Synergistically, when both domains are present in Kif14 molecules, they facilitate robust Kif14 tip tracking, which, we note, is not due to Kif14 super-processivity. Kif14 tip tracking is autonomous, and not dependent on auxiliary proteins, while other tip-tracking kinesins, like Ncd and MCAK, featuring the SxIP motif, depend on EB1 for localization to the microtubule tips [41, 42]. The only other example of a kinesin motor domain that preferentially binds to tubulin in its GTP over the GDP conformation and thus autonomously tracks the tips of growing microtubules is the non-motile ciliary kinesin Kif7 [43]. Kif14, thanks to its anchoring domain, is the only described kinesin that can both processively walk and autonomously microtubule tip-track.

Kif14 molecules, when comprising the diffusible domain and the motor domain, display two modes of interaction with microtubule lattices, directionally unbiased diffusion and unidirectional processive stepping towards the plus ends. These two modes of interaction exclude each other. We find that switching between them is regulated by dimerization. Monomeric Kif14 constructs, both in absence and in presence of the motor domain, diffuse on microtubules, while dimeric constructs, featuring motor domains, processively step. In part analogously, the constitutively dimeric kinesin-14 molecules, diffusing along microtubules via their tail anchoring domain, become activated upon binding of a second microtubule, to which they crosslink, and which they then actively slide [6, 7]. By contrast, Kif14 is activated, like other kinesin-3 family members [15, 44–46], by dimerization. Once activated, Kif14 molecules can walk processively as individual molecules.

To explain the enhanced processivity of the individual Kif14 molecules, we propose that the intrinsically disordered anchoring domain is a pliable tether for the Kif14 motor domains, keeping the kinesin bound to the microtubule in various configurations, even when one or motor domains are released (Fig. 6). This is similar to Kif1a, another kinesin-3, which can step processively as monomer, without detaching from the microtubule surface, due to a short, auxiliary, microtubule binding site, the K-loop, comprising less than two dozen amino acids, integral to the Kif1a motor-domains [47]. While processivity of monomeric Kif1a is caused by electrostatic interaction between the C-terminal tails of the microtubules and this motor-internal short loop [48], Kif14 interacts with microtubules electrostatically by means of the positively charged anchoring domain, which is distinct-from and external-to the motor domain. The anchoring domain is not required for processive Kif14 stepping. However, its presence greatly increases Kif14 run lengths and interaction times, rendering the kinesin super-processive. The kinesin-8 family members S. cerevisiae Kip3 and human Kif18A kinesin-8, similarly, are super-processive due to microtubule binding regions external of their motor domains [27, 28, 49–51]. Likewise, enhanced processivity could be mediated by an accessory protein binding to the motor and anchoring it to the microtubule, as is the case e.g. for S. cerevisiae kinesin-14 Kar3 [34, 52, 53] or kinesin-1, which can be anchored by the mitochondria adaptor protein TRAK [29].

**Fig. 6.**
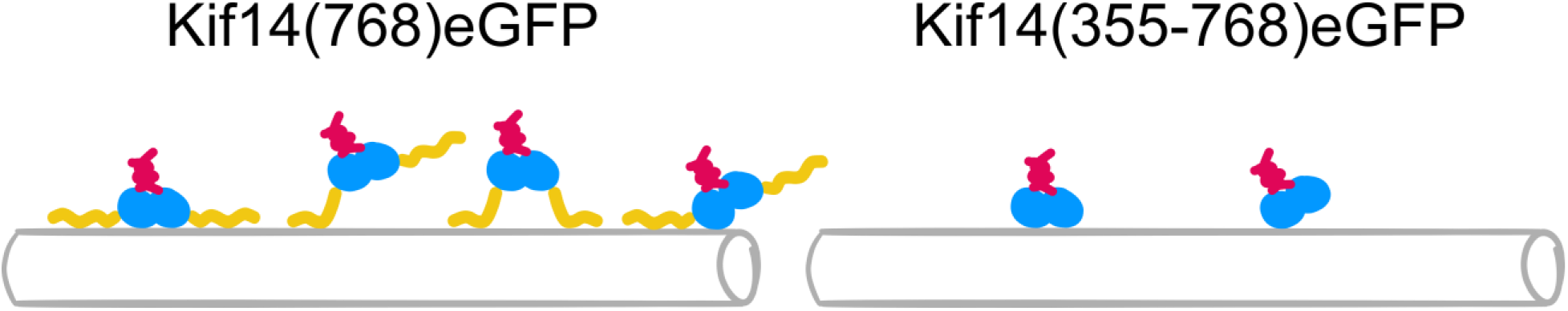
Schematic representation of various binding configurations of Kif14(768)eGFP (left) and Kif14(355-768)eGFP (right).

An important consequence of molecular motor anchoring and the accompanying increase in processivity is the enhancement of ability of the motor to move in crowded environments. Site-specific roadblocks and indistinct protein crowding decrease the run lengths of kinesins [54, 55]. In particular, enclosing microtubules with cohesive patches of intrinsically disordered microtubule-associated proteins like tau can prevent kinesin stepping in the covered microtubule regions altogether [30]. Microtubule-binding accessory proteins increase the processivity of kinesins allowing them to move efficiently in tau-covered regions [29]. Kif14 anchoring fulfills a similar role, holding the motor domains on a tether, which increases their processivity, and thus allows Kif14 pass through cohesive patches of tau molecules. As the midbody of the microtubule spindle, which forms during cell division, is especially crowded with proteins, we speculate that the Kif14 anchoring domain facilitates stepping in this highly crowded environment. Association with the microtubule crosslinker PRC1 in the midbody [17] might further promote the Kif14 stepping by providing an additional anchor. In vitro, Kif14 exhibits bimodal motion with diffusive monomeric and super-processive dimeric molecules, which make up 89% and 11% of the observed molecules respectively, but this ratio might be significantly shifted by the Kif14 interaction with microtubule-associated binding partners like PRC1.

In vivo, posttranslational modification, for example phosphorylation may regulate the functioning of the intrinsically disordered anchoring domain containing 39 phosphorylation sites as predicted by PHOSIDA [56], which will influence its binding to microtubules. In turn, posttranslational modifications of the microtubules themselves might also further regulate Kif14 stepping, diffusion and tip tracking. During cytokinesis, Kif14 is positioned between the overlapping microtubules of the spindle midzone. We here find Kif14 is a multi-versatile motor ideally equipped to locate-to and stabilize the overlapping microtubules at the spindle midzone (a) by stepping super-processively to microtubule plus ends, thus accumulating at the ends of the microtubules, without detaching from them (b) by binding preferentially the GTP-tubulin present at microtubule plus-ends and (c) by stably crosslinking parallel microtubules, while sliding antiparallel ones. All three functions, stepping, plus-end tracking and sliding/crosslinking, are promoted by the intrinsically disordered Kif14 anchoring domain, which tethers kinesin-3 to the microtubule lattice, exemplifying pliable protein tethering as fundamental mechanism of molecular motors regulation, complementary to ligand-binding and biochemical modification.

## METHODS

### PROTEIN EXPRESSION AND PURIFICATION

#### Homo sapiens

Kif14 constructs were obtained by PCR amplification of the amino acids 1-768 for Kif14(768)-eGFP, 355-768 for Kif14(355-768)-eGFP, 1-709 for Kif14(709)-eGFP, 1-355 for Kif14(355)-eGFP, and 1-715 for Kif14(715)-eGFP using primers containing AscI- and NotI-digestion sites flagging the particular Kif14-encoding nucleotide sequences. After AscI-NotI-digestion of the inserts, they were ligated into an AscI-NotI-digested pOCC destination vector containing an N-terminal MBP followed by a 3C PreScission Protease cleavage site and a C-terminal GFP tag followed by a 3C PreScission Protease cleavage site and a 6xHis-tag.

#### Homo sapiens

Kif5b truncated construct was obtained by PCR amplification of the amino acids 1-905 using primers containing AscI- and NotI-digestion sites flagging the particular Kif5b-encoding nucleotide sequences. After AscI-NotI-digestion of the inserts, they were ligated into an AscI-NotI-digested pOCC destination vector containing a C-terminal GFP tag followed by a 3C PreScission Protease cleavage site and a 6xHis-tag.

All proteins were expressed in SF9 insect cells using the opensource FlexiBAC baculovirus vector system for protein expression [57]. The insect cells were harvested after 4 days by centrifugation at 300 g for 10 min at 4 °C in an Avanti J-26S ultracentrifuge (JLA-9.1000 rotor, Beckman Coulter). The cell pellet was resuspended in 5 ml ice-cold phosphate buffered saline (PBS) and stored at - 80 °C for further use. For cell lysis, the insect cells were homogenized in 30 ml ice-cold His-Trap buffer (50 mM Na-phosphate buffer, pH 7.5, 5% glycerol, 300 mM KCl, 1 mM MgCl2, 0.1% tween-20, 10 mM BME, 0.1 mM ATP) supplemented with 30 mM imidazole, Protease Inhibitor Cocktail (cOmplete, EDTA free, Roche) and benzonase to the final concentration of 25 units/ml, and centrifuged at 45000 x g for 60 min at 4 °C in the Avanti J-26S ultracentrifuge (JA-30.50Ti rotor, Beckman Coulter). The cleared cell lysate was incubated in a lysis buffer-equilibrated Ni-NTA column (HisPur Ni-NTA Superflow Agarose, Pierce, VWR) for 2 h at 4 °C. The Ni-NTA column was washed with a wash buffer (His-Trap buffer supplemented with 60 mM imidazole) and the protein was eluted with an elution buffer (His-Trap buffer supplemented with 300 mM Imidazole). The fractions containing the protein of interest were pooled, diluted 1:10 in the His-Trap buffer and the purification tag was cleaved overnight with 3C PreScisson protease. The solution was reloaded onto a Ni-NTA column to further separate the cleaved protein from the 6xHis-tag. The protein was concentrated using an Amicon ultracentrifuge filter and flash frozen in liquid nitrogen. Kif14 constructs were finally separated from the MBP tag by size-exclusion chromatography on a superpose 6 10/300 (GE Healthcare) column equilibrated with BRB80 buffer (80 mM Pipes (pH 6.8), 2 mM MgCl_2_, 1 mM EGTA).

### MICROTUBULE PREPARATIONS

Tubulin was isolated from pig brains and labeled as described previously [58–60].

For preparation of biotinylated microtubules isolated tubulin was mixed with biotinylated tubulin (Cytoskeleton Inc., T333P) at 50:1 mass ratio. For preparation of rhodamine-labeled microtubules isolated tubulin was mixed with rhodamine-labeled tubulin (Cytoskeleton Inc., TL620M) at 5:1 mass ratio. For preparation of Alexa647-labeled microtubules isolated tubulin was mixed with Alexa647-labeled tubulin at 20:1 mass ratio.

GMPCPP-stabilized microtubules were grown using a mixture of 1.3 mg ml^−1^ tubulin and 2 mM GMPCPP (Merck, NU-405L) in BRB80 and incubated for 1 hour (stepping and photobleaching assay) or 15 min (seeds for dynamic microtubule assay and transport microtubule for sliding assay) at 37°C. The mixture was then spun at 14000 g in a tabletop centrifuge (Beckman Coulter, Microfuge 18). Finally, the supernatant was discarded, and the pellet was resuspended in 50 μl BRB80.

### TIRF IMAGING

Imaging was carried out at room temperature using an inverted microscope (Nikon, Ti-E Eclipse) equipped with a 100x or 60x 1.49 N.A. oil immersion objective (Nikon, Plan Apo) and an EMCCD camera (Andor Technology, iXon Ultra 888). Samples were excited using LU-N4/N4S laser unit (Nikon). Illumination and image acquisition was controlled by NIS Elements Advanced Research software (Nikon). Unlabeled microtubules and GFP-labeled proteins were visualized sequentially by switching between 488 nm laser in TIRF mode and interference reflection microscopy (IRM) mode [31] and the corresponding filter set. Acquisition rates ranged from one frame per 0.12 s to one frame per 10 s, and are indicated as time scale bars in the individual figures.

### IN VITRO ASSAYS

Flow chambers for TIRF imaging assays were prepared as described previously [61, 62]. Channels were treated with anti-biotin (Sigma, B3640, 1 mg ml^−1^ in PBS) or anti-β-tubulin (Sigma, T7816, 1 mg ml^−1^ in PBS) antibody solution for 5 minutes, followed by one-hour incubation with 1% Pluronic F127 (Sigma, P2443).

For the stepping or photobleaching assays the channels were washed with 40 μl of buffer A (20 mM Hepes (pH 7.3), 75 mM KCl, 2 mM MgCl_2_, 1 mM EGTA). Subsequently Alexa647-labeled or biotinylated fluorescently unlabeled microtubules were injected into the channel. Then, the channels were flushed with 20 μl of buffer A to remove unbound microtubules. Finally, the flow chamber was suffused with a Kif14 construct diluted in the assay buffer (buffer A supplemented with 0.5 mg ml^−1^ casein, 10 mM dithiothreitol, 0.1% Tween-20, 20 mM D-glucose, 22.4 μg ml^−1^ glucose oxidase, and 20 μg ml^−1^ catalase), with or without 1 mM ATP as indicated in the main text.

For the dynamic microtubules assay channels were washed with 40 μl of BRB80 (80 mM Pipes (pH 6.8), 2 mM MgCl_2_, 1 mM EGTA). Subsequently biotinylated fluorescently unlabeled microtubules were injected into the channel. Then, the channels were flushed with 20 μl BRB80 to remove unbound microtubules. Finally, the flow chamber was suffused with a mixture of 20 µM soluble tubulin and a Kif14 construct in polymerization buffer (BRB80 supplemented with 20 mM D-glucose, 22.4 μg ml^−1^ glucose oxidase, 20 μg ml^−1^ catalase, 1 mM ATP and 1 mM GTP).

For microtubule sliding assay channels were washed with 40 μl of buffer A with subsequent injection biotinylated unlabeled microtubules. Then, the channels were flushed with 20 μl buffer A to remove unbound microtubules. Finally, the flow chamber was suffused with a mixture of a Kif14 construct with rhodamine-labeled transport microtubules diluted in assay buffer with 1 mM ATP.

For the estimation of binding affinities, 40 μl of buffer A with subsequent injection of unlabeled microtubules. Then, the channels were flushed with 20 μl buffer A to remove unbound microtubules. Next, the flow chamber was suffused with solutions of a GFP-tagged Kif14 construct at increasing concentrations, in assay buffer with or without 1 mM AMPPNP or ADP. Several snapshots were taken for each concentration using IRM to note the position of the microtubules and using the 488 nm channel to quantify the amount of microtubule-bound motor.

### IMAGE ANALYSIS

Tracking of single Kif14(768)-eGFP molecules for the estimation of diffusion coefficient was performed using FIESTA [63] software. Processive Kif14(768)-eGFP molecules were distinguished from diffusive ones using kymographs in FIESTA. A molecule was considered processive if it exhibited continuous unidirectional movement along a microtubule with back steps no larger than 2 pixels. The diffusive traces were then used to estimate mean square displacement (MSD). Next, the MSD values were fitted using the MATLAB curve-fitting toolbox (linear fit, MSD = 2Dτ + C, D - diffusion coefficient, τ - lag time, C - offset).

Velocities, run lengths of Kif14(768)-eGFP and Kif14(355-768)-eGFP, microtubule sliding velocities, and dynamic microtubule parameters were analyzed manually in FIJI v.1.52 [64]. Distribution of run lengths was estimated using Kaplan-Meier survivor function in Matlab (Mathworks) [65]. Kymographs were generated using KymographBuilder plugin in FIJI. Intensity profiles were generated using ‘Plot Profile’ function in FIJI.

Photobleaching evaluation was performed using FIESTA. The positions of bleaching steps were first estimated by eye. For each molecule initial intensity was calculated by averaging all intensities before the first bleaching step.

The analysis of the images for the estimation of binding affinities was performed manually in FIJI v.1.52. The background subtracted intensity of GFP signal along microtubules was taken as a readout of a Kif14 construct binding to microtubules. The Kd values were calculated by fitting the data points to the Hill equation using Matlab.

### BIOINFORMATICS ANALYSIS

Disorder degree was analyzed using IUPred2 and ANCHOR2 interfaces [66–69]. Coiled-coil prediction was performed in PCOILS (MPI bioinformatics toolkit) [70–72]. Net charge per residue and the isoelectric point were estimated by localCIDER [73].

## ACKNOWLEDGEMENTS

We thank Radan Matura, Verena Henrichs and Lenka Grycova for help with protein preparation, Valerie Siahaan for help with preparation of tau islands, and Yulia Bobrova for technical support. We acknowledge the financial support from the Czech Science Foundation (grant no. 19-27477X to Z.L. and 20-04068S to M.B.) and the Charles University Grant Agency (grant no. 1372218 to I.Z.), the Introduction of New Research Methods to BIOCEV (CZ.1.05/2.1.00/19.0390) project from the ERDF, the institutional support from the CAS (RVO: 86652036) and the Imaging Methods Core Facility at BIOCEV, an institution supported by the Czech-BioImaging large RI projects (LM2015062 and CZ.02.1.01/0.0/0.0/16_013/0001775, funded by MEYS CR) for their support in obtaining imaging data presented in this paper.

## AUTHOR CONTRIBUTION

Conceptualization, M.B., Z.L.; Methodology, I.Z., M.B., Z.L.; Formal Analysis, I.Z.; Investigation, I.Z.; Resources, I.Z., S.D.; Writing, I.Z., M.B., Z.L.; Visualization, I.Z.; Supervision, M.B., Z.L.; Funding Acquisition, I.Z., M.B., Z.L.

---

**Fig. S1.**
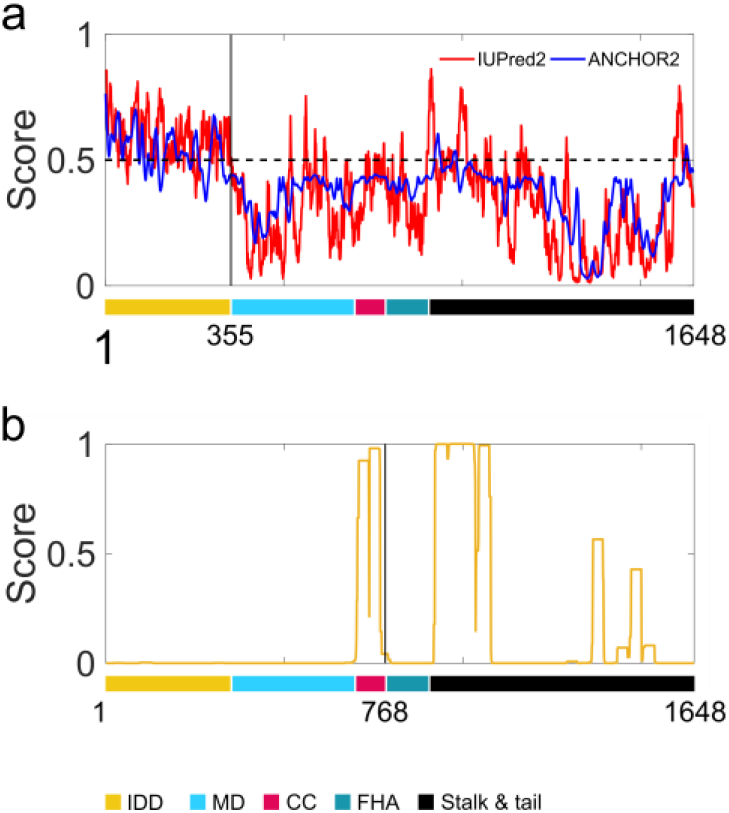
Bioinformatic analysis of full-length Kif14. **(a)** Analysis of the disorder degree by IUPred2 (red) and ANCHOR2 (blue) interfaces. A given region is considered disordered when the disorder probability is above 0.5.; bold vertical line shows the end of the intrinsically disordered N-terminal domain **(b)** Coiled-coil prediction by PCOILS. Bold vertical line shows the C-terminus of Kif14(768)-eGFP construct.

**Fig. S2.**
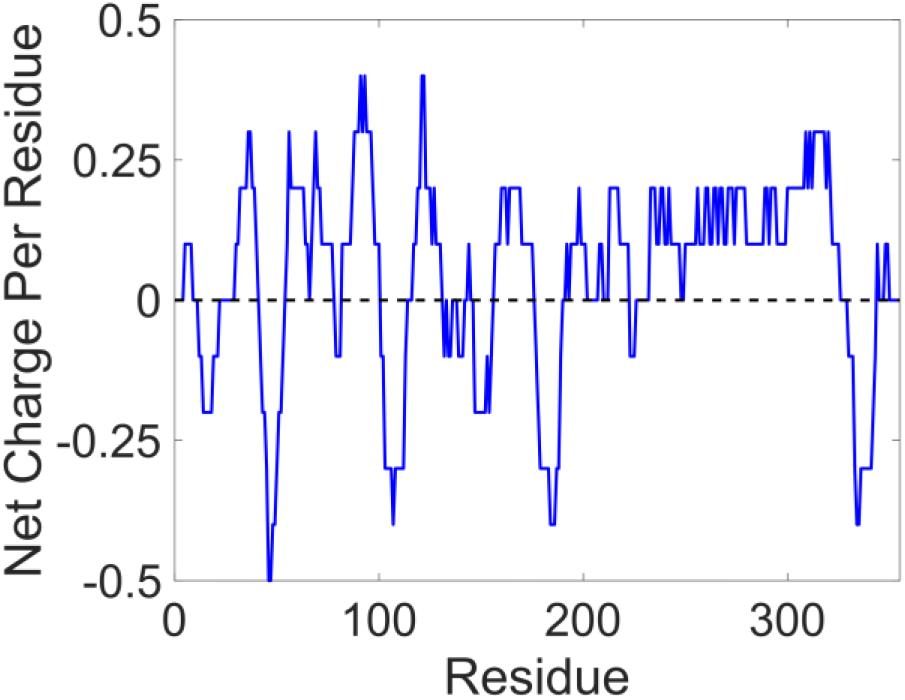
Net charge per residue prediction of the N-terminal (1-355 aa) domain of Kif14.

**Fig. S3.**
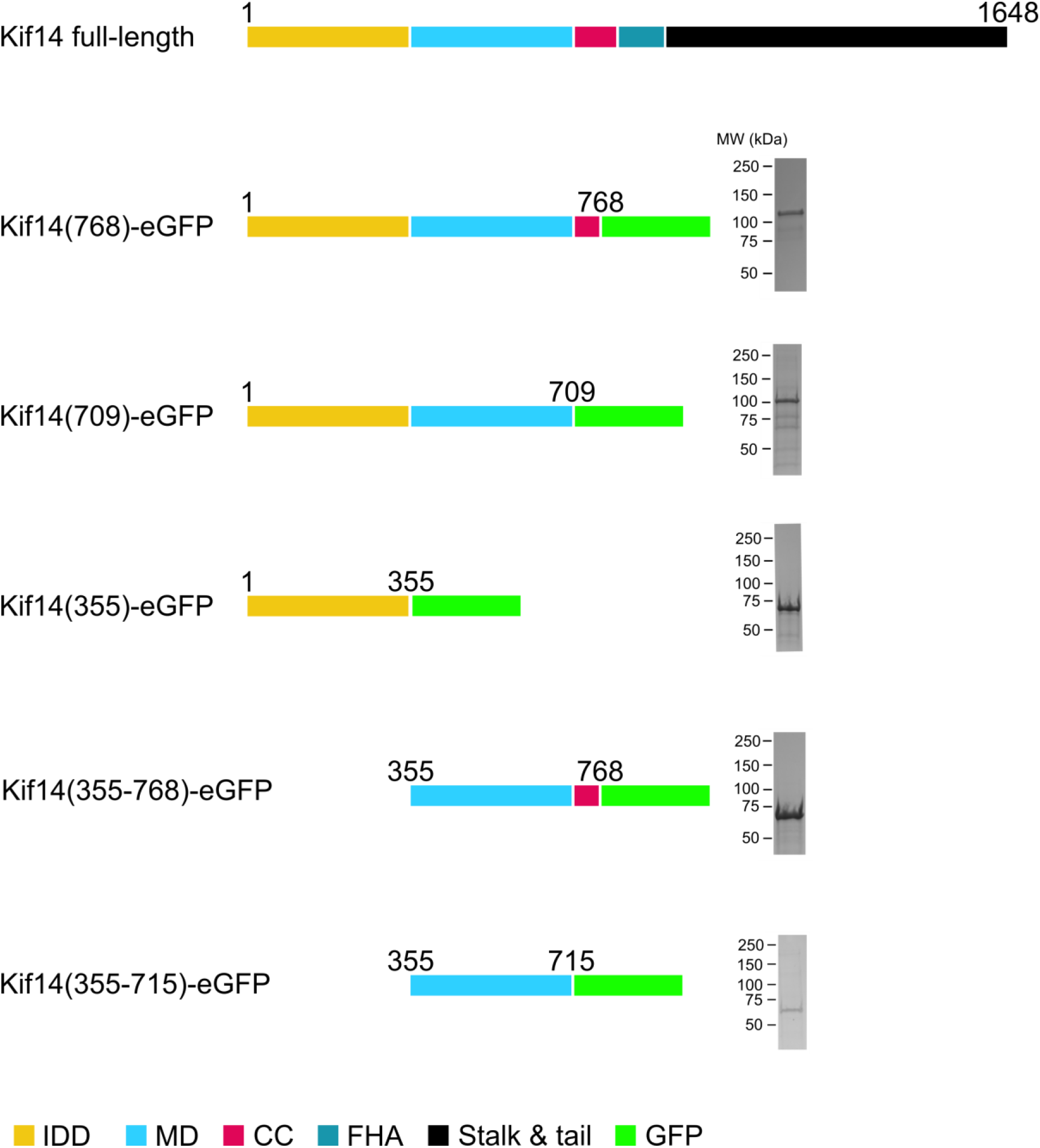
Schematic diagrams of the GFP-labeled Kif14 constructs with corresponding coomassie-stained SDS-polyacrylamide gel electrophoresis (SDS-PAGE) of the purified recombinant proteins. The full-length Kif14 consists of the N-terminal intrinsically disordered domain (aa 1-355), the motor domain (MD, aa 355-709), a coiled-coil (CC, aa 709-800), a forkhead associated domain (FHA, aa 800-900), consecutive stalk and tail (aa 900-1648).

**Fig. S4.**
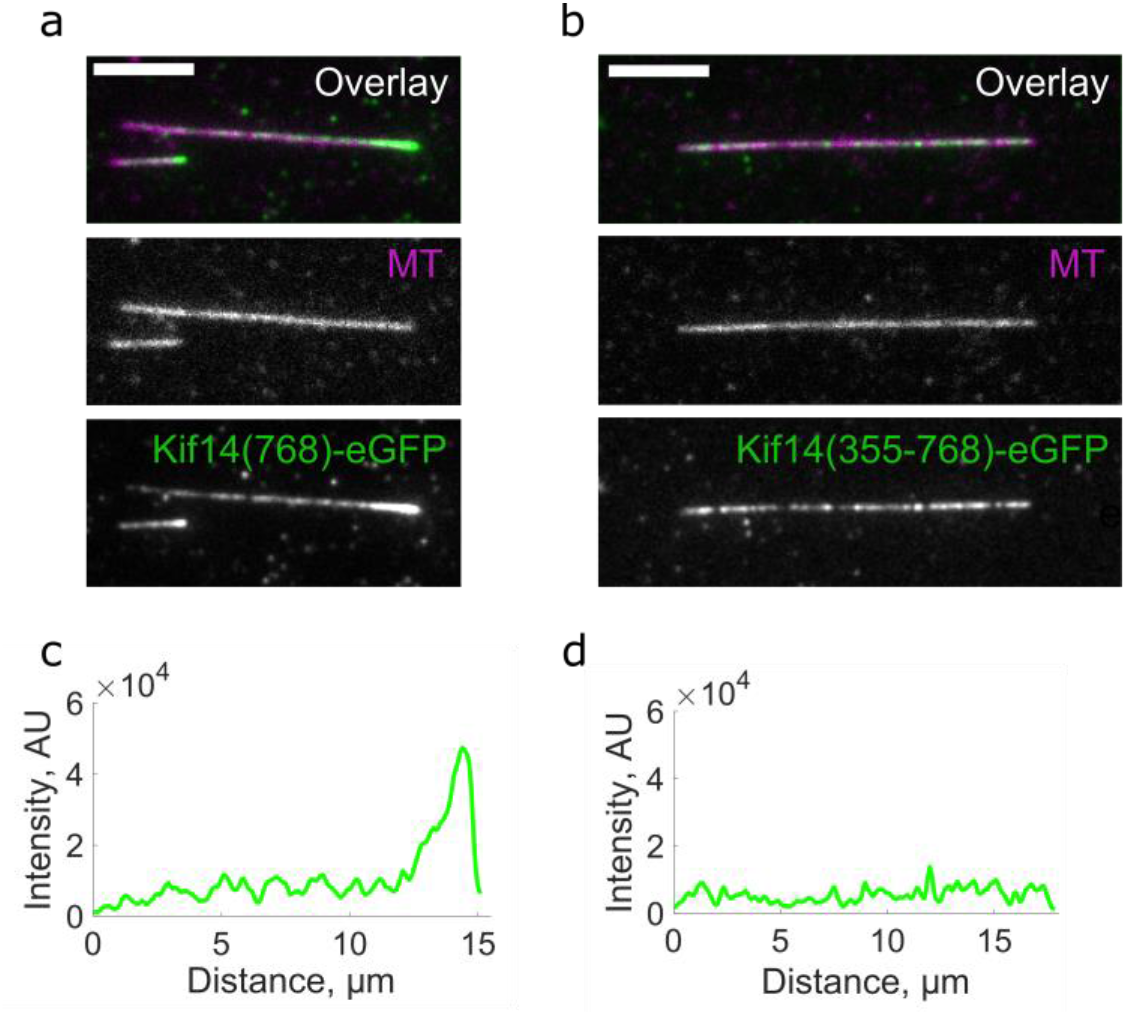
The intrinsically disordered domain promotes microtubule plus end accumulation of Kif14. Snapshots of **(a)** 1 nM Kif14(768)-eGFP and **(b)** 25 nM Kif14(355-768)-eGFP decorating Alexa647-labeled microtubule after 5 min incubation, scale bars: 5 μm. **(c)** and **(d)** Intensity distribution of the GFP signal along the microtubules in (a) and (b), respectively.

**Fig. S5.**
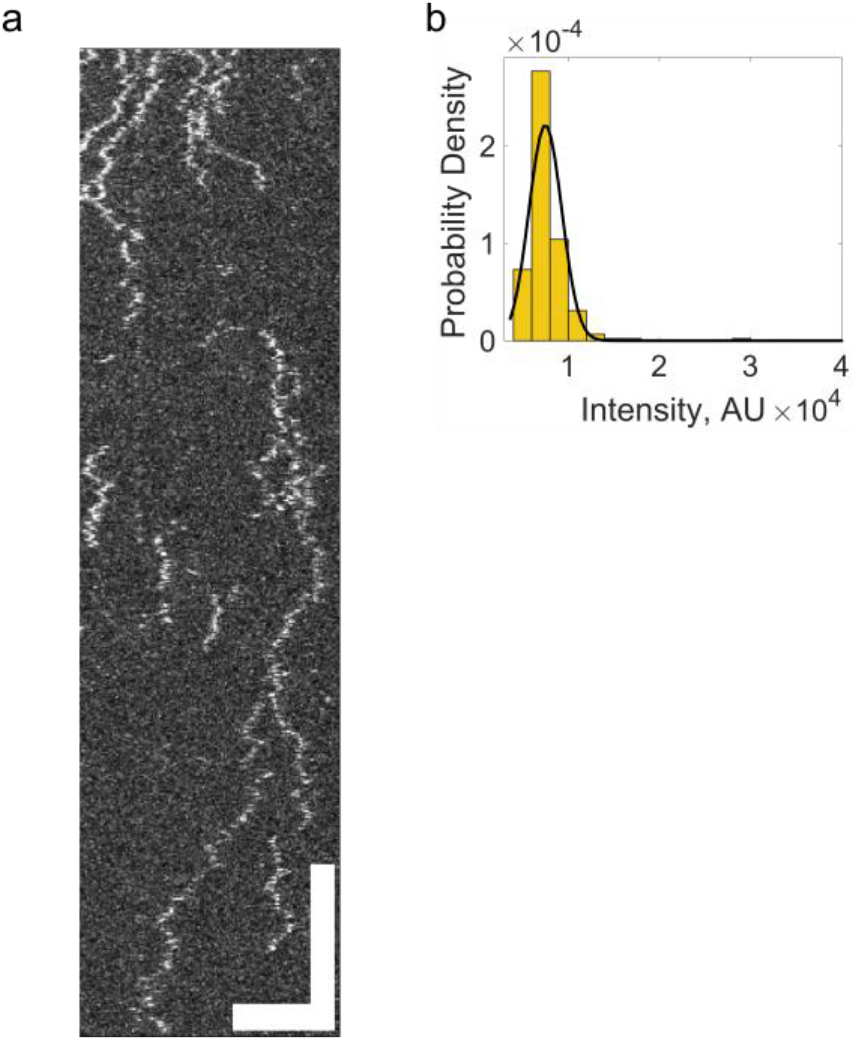
Evaluation of the constitutively monomeric Kif14(709)-eGFP construct. **(a)** Kymographs representing interaction of Kif14(709)-eGFP with microtubules (unlabeled). Horizontal scale bar 5 μm; vertical bar 20 s. (**b**) GFP signal intensity distribution of Kif14(709)-eGFP in the nucleotide-free state. Black lines indicate Gaussian fits to the intensity histograms.

**Fig. S6.**
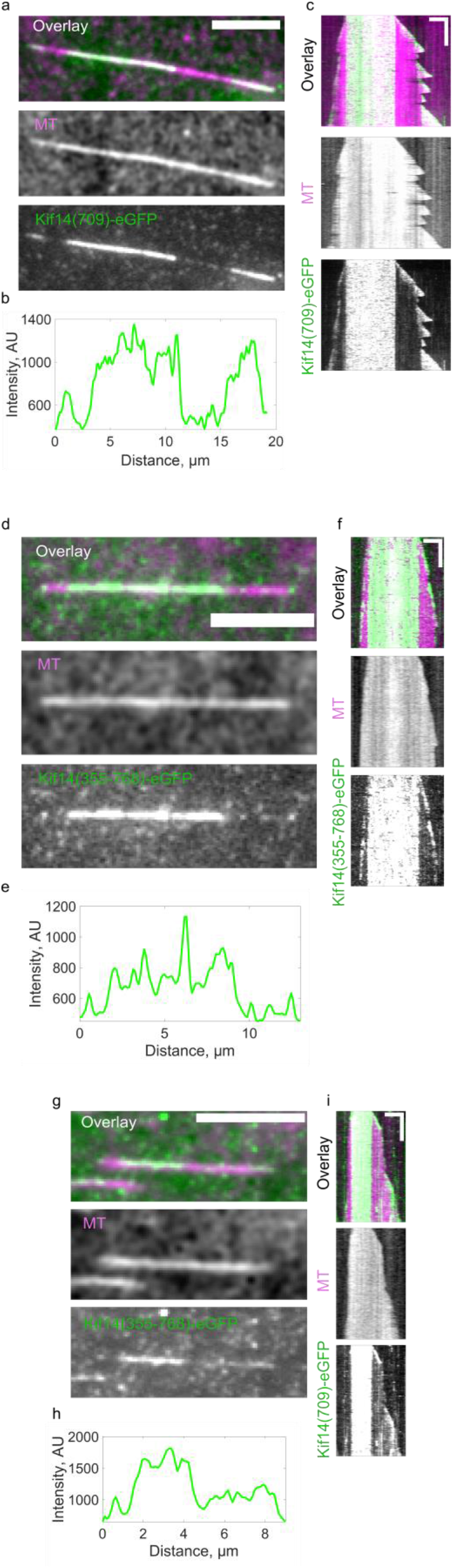
Kif14 constructs autonomously track the tips of growing microtubules. **(a)** Snapshots of a dynamic microtubule (magenta) in the presence of 140 nM Kif14(709)-eGFP (green). **(b)** Intensity distribution of the GFP signal along the microtubules in (a). **(c)** Kymographs representing 140 nM Kif14(709)-eGFP (green) interaction with the polymerizing microtubule (magenta). Horizontal scale bar 3 μm; vertical bar 5 min. **(d)** Snapshots of a dynamic microtubule (magenta) in the presence of 150 nM Kif14(355-768)-eGFP (green). **(e)** Intensity distribution of the GFP signal along the microtubules in (d). **(f)** Kymographs representing 150 nM Kif14(355-768)-eGFP (green) interaction with the polymerizing microtubule (magenta). Horizontal scale bar 3 μm; vertical bar 5 min. **(g)** Snapshots of a dynamic microtubule (magenta) in the presence of 2.5 µM Kif14(355)-eGFP (green). **(h)** Intensity distribution of the GFP signal along the microtubules in (g). **(i)** Kymographs representing 2.5 µM Kif14(355)-eGFP (green) interaction with the polymerizing microtubule (magenta). Horizontal scale bar 3 μm; vertical bar 5 min.

**Fig. S7.**
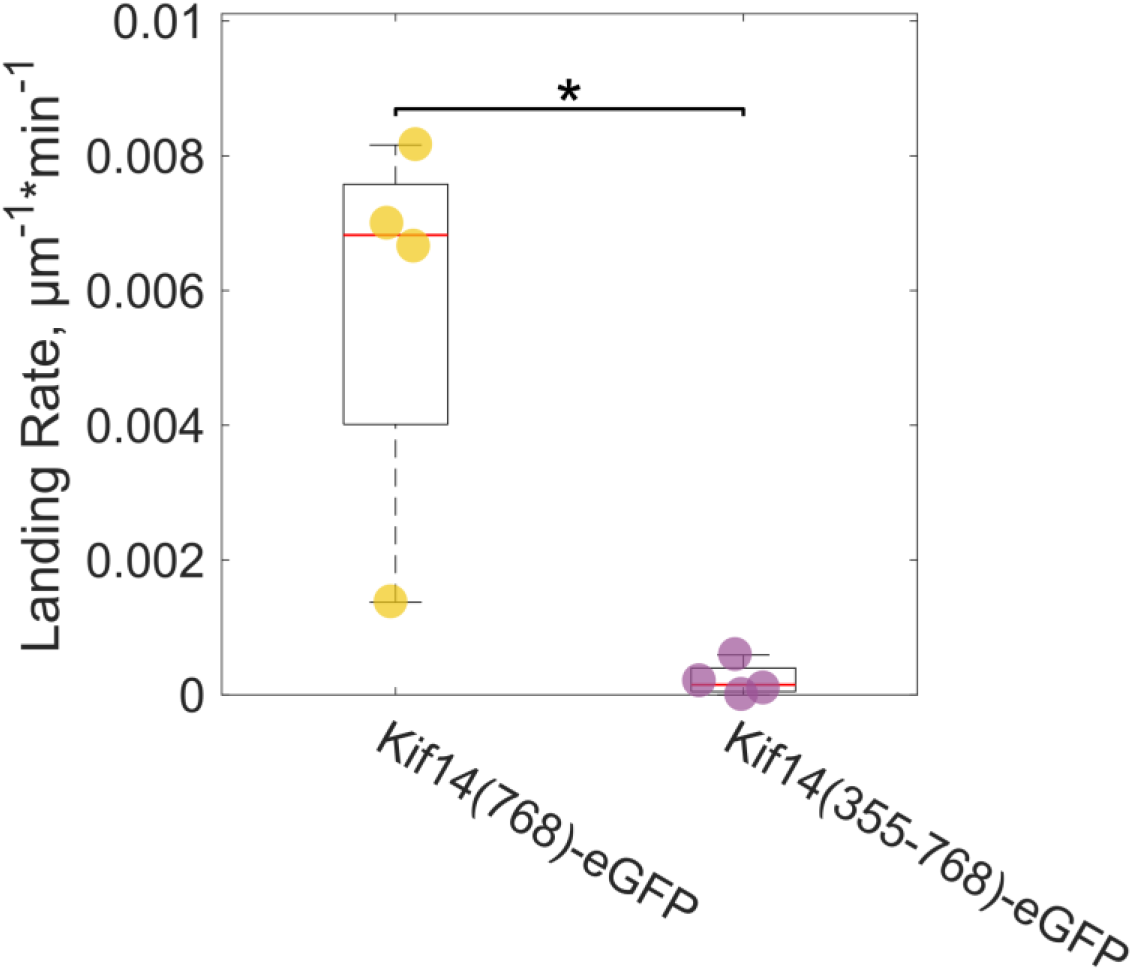
Landing rate of transport microtubules on template microtubules in the presence of 20 nM Kif14 constructs. *p < 0.05, determined by Student t test

**Fig. S8.**
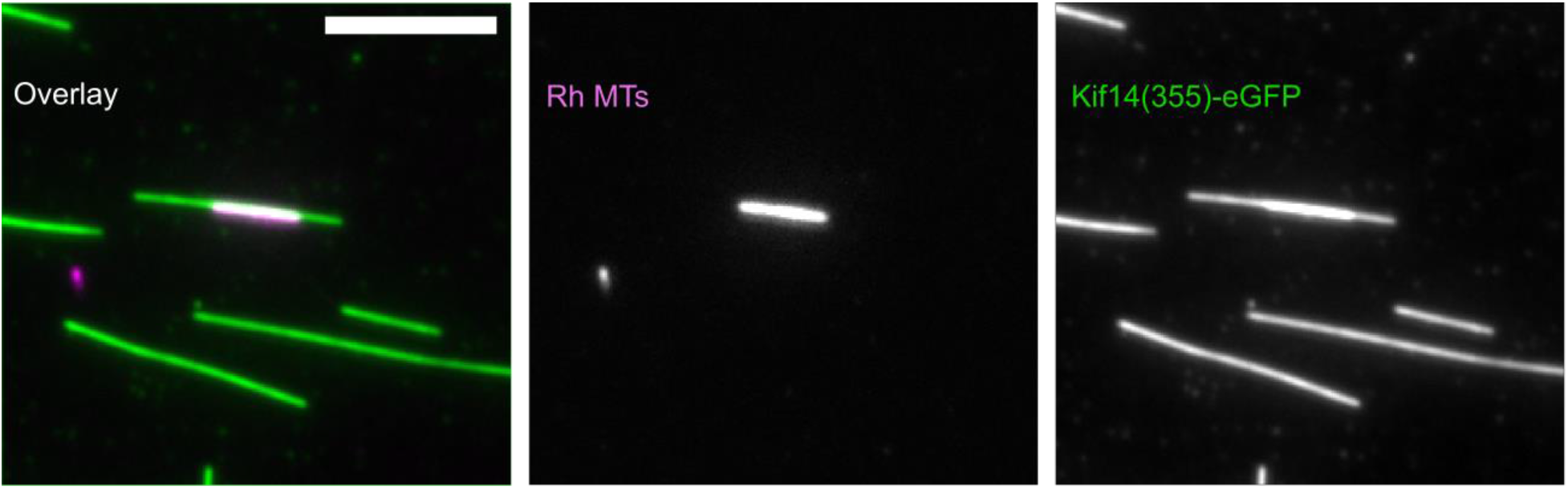
Fluorescence micrographs of Kif14(355)-eGFP (green) crosslinking microtubules: template microtubules (unlabeled) and transport microtubules (magenta). Scale bar, 5 μm.

**Movie 1.** Kif14(768)-eGFP (green) accumulates at the plus end of the microtubules (magenta). Scale bar 5 μm.

**Movie 2.** Kif14(355-768)-eGFP (green) does not accumulate at the plus end of the microtubules (magenta). Scale bar 5 μm.

**Movie 3.** Movement of Kif14(768)-eGFP (green) on the microtubule (unlabeled) covered with tau-mCherry cohesive islands (magenta). Scale bar 3 μm.

**Movie 4.** Movement of Kif14(355-768)-eGFP (green) on the microtubule (unlabeled) covered with tau-mCherry cohesive islands (magenta). Scale bar 5 μm.

**Movie 5.** Dynamic microtubules (magenta) in the presence of Kif14(768)-eGFP (green). Scale bar 5 μm.

**Movie 6.** Dynamic microtubules (magenta) in the presence of Kif14(709)-eGFP (green). Scale bar 5 μm.

**Movie 7.** Dynamic microtubules (magenta) in the presence of Kif14(355)-eGFP (green). Scale bar 5 μm.

**Movie 8.** Dynamic microtubules (magenta) in the presence of Kif14(355-768)-eGFP (green). Scale bar 5 μm.

**Movie 9.** Rhodamine-labeled transport microtubule (magenta) sliding along a template microtubule (unlabeled) in the presence of Kif14(768)-eGFP. Scale bar 3 μm.

